# *When* makes you unique: temporality of the human brain fingerprint

**DOI:** 10.1101/2021.03.24.436733

**Authors:** Dimitri Van De Ville, Younes Farouj, Maria Giulia Preti, Raphaël Liégeois, Enrico Amico

## Abstract

The extraction of “fingerprints” from human brain connectivity data has become a new frontier in neuroscience. However, the time scales of human brain identifiability have not been addressed yet. In other words, what temporal features make our brains more “identifiable”? We here explore the dynamics of brain fingerprints (or brainprints) along two complementary axes: 1) *what is the optimal time scale* at which brainprints integrate sufficient information, 2) *when best* identification happens. Using dynamic identifiability, we show that the best identification emerges at longer time scales (~300*s*); however, short transient “bursts of identifiability” persist even when looking at shorter functional interactions. We find that these bursts of identifiability might be strongly associated with neuronal activity. Furthermore, we report evidence that different parts of connectome fingerprints relate to different time scales: i.e., more visual-somatomotor at short temporal windows, more frontoparietal-DMN driven by increasing temporal windows. Finally, using a meta-analytic approach, we show that there is a broad spectrum of associations between brainprints and behavior. At faster time scales, human brain fingerprints are linked to multisensory stimulation, eye movements, affective processing, visuospatial attention. At slower time scales instead, we find higher-cognitive functions, such as language and verbal semantics, awareness, declarative and working memory, social cognition. We hope that this first investigation of the temporality of the human brain fingerprint will pave the way towards a better understanding of *what* and *when* makes our brains unique.

## Introduction

In the 17th century, physician Marcello Malpighi observed the existence of distinctive patterns of ridges and sweat glands on fingertips ^1^. This was a major breakthrough, and originated a long and continuing quest for ways to uniquely identify individuals based on fingerprints, a technique massively used until today. In the modern era, the concept of fingerprinting has expanded into other sources of data, such as voice recognition and retinal scans, among many others ^2^. It is only in the past few years that technologies and methodologies have achieved high-quality measures of an individual’s brain to the extent that personality traits and behavior can be characterized. The most insightful correlates emerged from the investigation of functional and structural connectivity that can be modeled and analyzed using network science. This area of research is usually referred to as Brain Connectomics^3^.

In Brain Connectomics, network organization of the brain is studied using either the structural or functional connectome. For the former, strength of white-matter pathways between pairs of brain regions is extracted from diffusion weighted imaging (DWI) data and referred to as structural connectivity (SC). For the latter, temporal statistical dependencies between pairs of activity time courses taken from functional magnetic resonance imaging (fMRI) define functional connectivity (FC). The most common paradigm for FC is resting-state fMRI during which subjects in the scanner are not engaging in a particular task^4^.

The concept of “fingerprints of the brain” is very novel ^5,6^ and has been boosted thanks to a seminal publication by Finn et al. ^5^ in 2015. They were among the firsts to show that, to a great extent, it is possible to robustly identify the functional connectome of a “target” subject from a sample database of FCs, simply by computing the spatial (Pearson) correlation of the target FC against the database ones. The success rate of this identification procedure based on brain connectivity data was above 90% for resting-state sessions, and ranged between 54% and 87% when including task-task and task-rest sessions ^5^. This seminal work demonstrated that an individual’s functional brain connectivity profile is both unique and reliable, similarly to a fingerprint, and that it is possible, with near-perfect accuracy in many cases, to identify an individual among a large group of subjects solely on the basis of her or his connectivity profile. These findings further incentivized human neuroimaging studies to move from inferences at the population level to results that apply to the single-subject level; i.e., by examining how individuals’ networks are functionally organized in unique ways ^7,8^, or by relating this functional organization to behavioral phenotypes in both health and disease ^9^, or even by implementing ways for maximizing and denoising fingerprints in brain data ^10^.

Yet, the discovery of brain fingerprints opened up a plethora of new questions, in particular, what exactly is the information encoded in brain connectomes that ultimately leads to correctly differentiating someone’s connectome from anybody else’s? In other words, what makes our brains unique? More specifically, related to the temporality of FC-based fingerprinting, is the brain more unique at some moments, and what is the temporal extent needed for a fingerprint to unfold?

Here we address these questions by tapping into the temporal dynamics of human brain connectivity. We use dynamic functional connectivity techniques to explore the time scales of brain identification; i.e., when and over which duration do these unique fingerprints originate, and which brain areas are most responsible for this. We demonstrate that optimal fingerprints manifest at a time scale of 300 sec based on dynamic functional connectomes. Nonetheless, unique individual “snapshots” of brain connectivity emerge at much shorter time scales already. In addition, snapshots at different time scales reveal specific connectivity patterns in terms of regions and functional networks, which shows how fMRI BOLD fluctuations relate to different types of underlying neuronal events. Moreover, when looking at different areas in the brain fingerprints, we noticed that subcortical regions are the fastest ones for individual identification; visual and somatomotor regions appear right after; ultimately, at slower time scales, frontoparietal and DMN emerge. Finally, a meta-analytical investigation revealed that brain fingerprints can be associated with behaviorally-relevant arrangement, revealing a complex gradient of relationship between the time scale of fingerprinting and digression from sensory behavioral traits to higher-order cognitive functions. In sum, for the first time, dynamic FC methods allowed us to investigate the temporality of brain fingerprints. We provide evidence that what makes our brains unique is multifaceted, based on *when and how long*. That is, individual identification is a temporally integrating and fluctuating feature of brain fingerprints.

## Results

We introduce “dynamic brain fingerprints” (or *brainprints*) to investigate the temporality of brain fingerprinting. The general scheme for dynamic identification can be divided in three steps (Fig. 1): 1) the time scale is set by the choice of the temporal window length; 2) sliding-window dynamic functional connectome frames (dFCs) are computed for each window position; 3) the similarity of frames within-subjects and between-subjects is evaluated, with the aim to extract the best “identifiable” connectome frames, for each subject (Fig. 1, see also Supplementary Fig. S1). In a nutshell, the temporal exploration of human brainprints can be decomposed according to two complementary axes: the *time scale* of brain identification or how long it takes for the information to optimally integrate, and the *best matching time* of identification or when best information is available (Fig. 1). This concept can be formally encoded into a “dynamic Identifiability matrix” (Fig. S1), in which the blocks represent within-subject dFCs similarity, and off block-diagonal elements contain the information on the between-subjects dFCs similarity (Fig. S1, see also Methods for details).

**Fig. 1.**
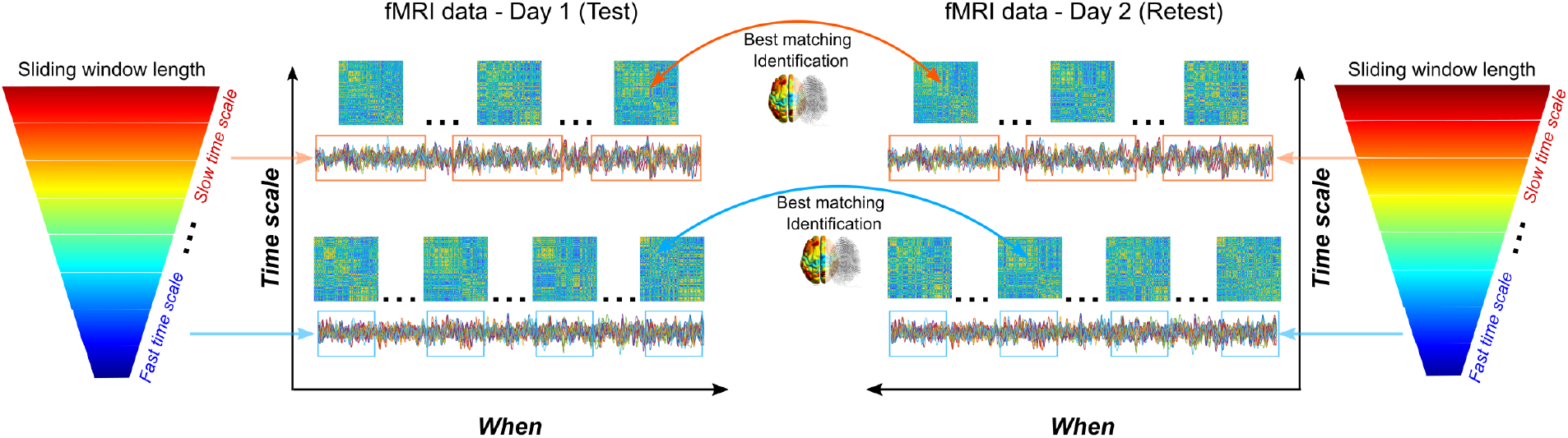
Exploring dynamic brain fingerprints. Schematic of dynamic connectome identification, for one subject. First, the *time scale* (window length) of the exploration, here depicted as a gradient cone, is set; secondly, dynamic functional connectivity frames are computed at each window, for both test and retest fMRI data; finally, the best matching frames across test and retest data are retrieved for identification.

We explored these temporality aspects of human brainprints across different time scales (window lengths) on the 100 unrelated subjects of the Human Connectome Project dataset. We started by selecting six different window lengths (7.2s, 36s, 72s, 144s, 288s, 576s, with a fixed sliding window step of 7.2 seconds), and explored dynamic differential identifiability (*d*Idiff) at every window (Fig. 2, see also Methods for details on the implementation).

**Fig. 2.**
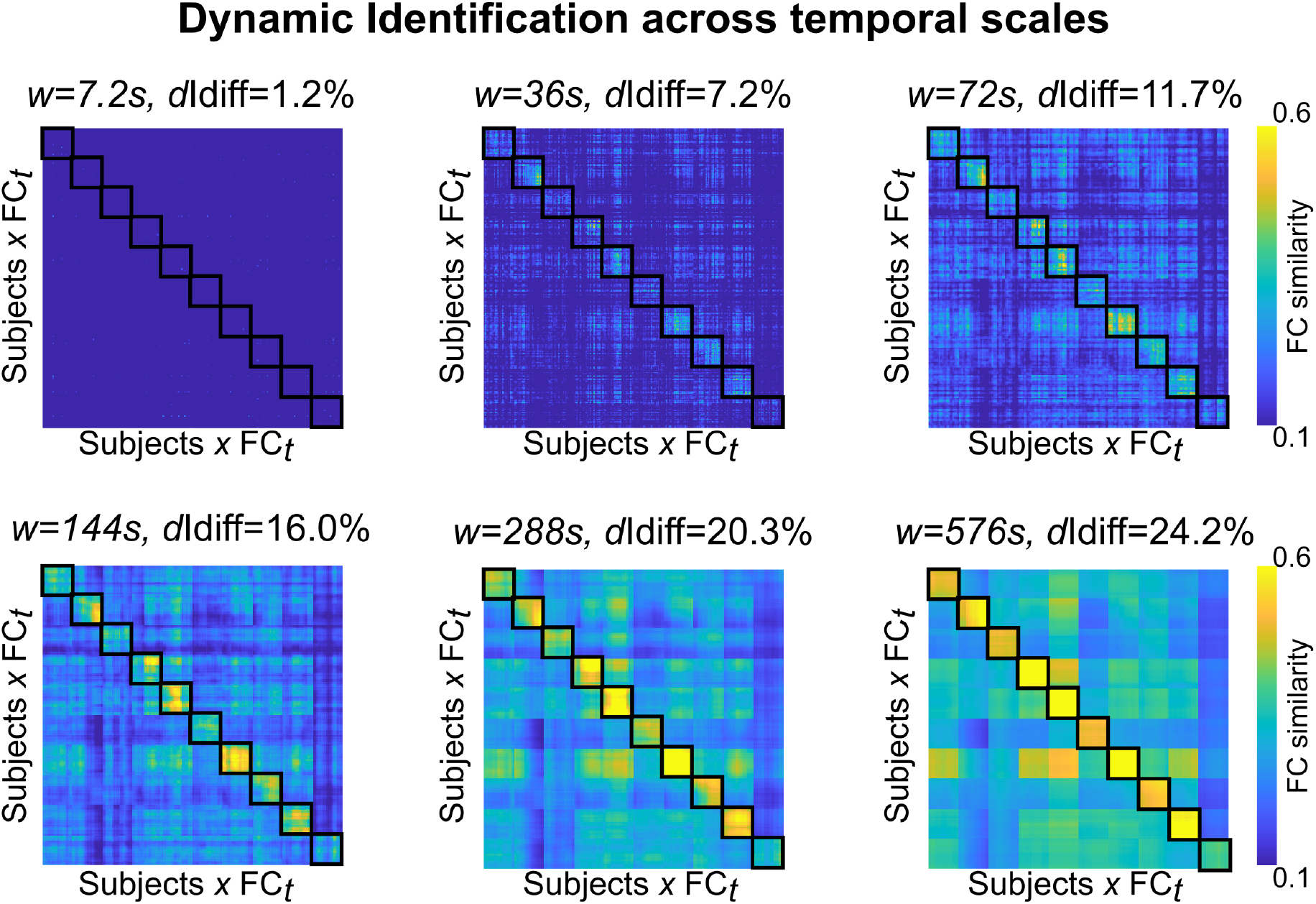
Dynamic Identification across temporal scales. Dynamic Identifiability matrix evaluation at six different window lengths: (7.2s, 36s, 72s, 144s, 288s, 576s). To ease its visualization, the dynamic identifiability matrix in the figure was reduced to 10 sample subjects. The dynamic differential identifiability on top of each matrix provides a score of the fingerprint level of the dataset across temporal scales. Black squares indicate the self similarity of each subject’s dFC frames.

The dynamic differential identification increases steadily with longer window lengths (Fig. 2). This is expected since we rely upon more time points for dFCs computation, increasing the stability of the functional connectivity profiles across test-retest sessions. However, clearly noticeable diagonal blocks respecting the subject boundaries start appearing before the maximal *d*Idiff window length (Fig. 2).

This suggests that, even at shorter time scales, there exist specific *brainprints* able to reliably link test-retest sessions.

We then explored if there were specific individual dFC frames that would be driving the dynamic identification. We therefore ranked the frames based on how good they could represent the subject (or *d*Iself, see Methods for details) across test-retest, and evaluated how good these dFC frames could also separate between subjects, via *d*Iothers. Fig. 3 shows that few FC frames can drive identifiability, especially at shorter window length (steeper curves, Fig. 3A). Note that, even if the *d*Iself behavior is expected since we based our dFC frames ranking on it, the fact that *d*Iothers (and consequently *d*Idiff) might follow the same trend is not trivial (Fig. 3A). Interestingly, when looking at the variability of the Top frame for identification across subjects, one can notice the emergence of characteristic spatial patterns of connectivity becoming more homogeneous in the population (Fig. 3B). Starting from visual and somatomotor patterns of connectivity at shorter temporal windows, degrading towards fronto-parietal network connectivity at slower time scales, finally including default mode network connectivity (Fig. 3B).

**Fig. 3.**
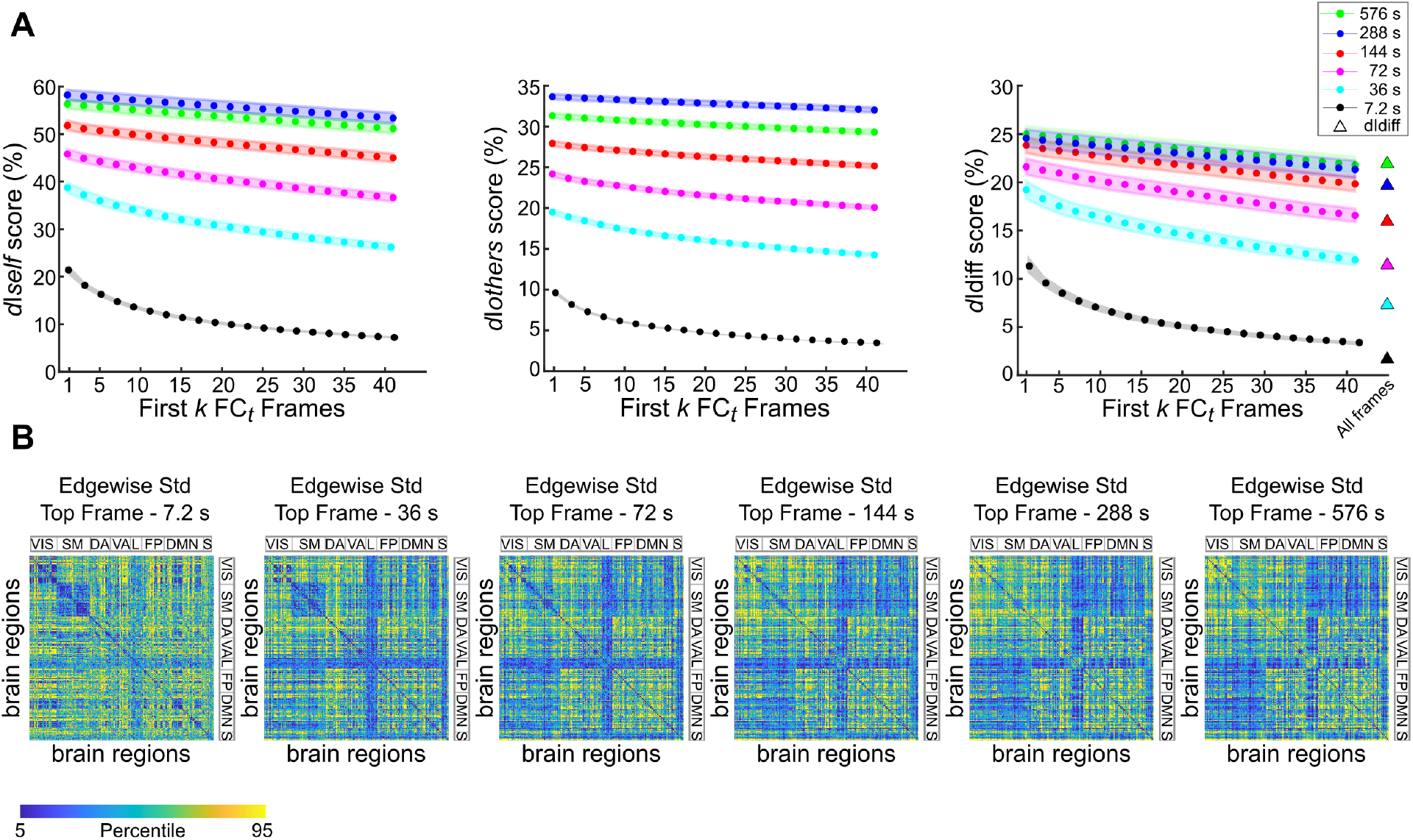
Brain fingerprint resides in few FC frames. **A)** Evaluation of brainprints across temporal scales through d*I*self (left), d*I*others (middle) and d*I*diff scores (right), when ranking dFC frames based on individual *d*Iself, in descending order. The *d*Idiff scores obtained with the ranking are compared with the ones obtained when taking all frames (triangles), also depicted in Fig. 2. **B)** Edgewise standard deviation across subjects of the best matching dFC frames, at each temporal scale. The matrices are ordered according to the seven resting state subnetwork organization proposed by Yeo and colleagues^12^, specifically: Visual (VIS), somatomotor (SM), dorsal attention (DA), ventral attention (VA), Limbic (L), Frontoparietal (FP) and default mode network (DMN). For completeness, an eight subcortical subnetwork (S) was added at the end (see Brain Atlas in Methods for details).

To evaluate the contribution of BOLD fluctuations that are most likely driven by neuronal activity-inducing signals, we repeated the same analysis using transient activity time courses (as in ^11^) obtained as the derivative of the deconvolved BOLD signals (Fig. S2). This simultaneously removes the effect of the hemodynamic response function (HRF) and detects transition moments in neural activity. The same finding as before is confirmed across *d*Iself, dIothers and *d*Idiff, with small fluctuations across temporal scales (Fig. S2).

Given the observed spatial variation as a function of time scale, we investigated in more details the spatial profiles of the fingerprints. Specifically, we used edgewise intra-class correlation (ICC, see Methods) to explore the FC connections most contributing to brain fingerprinting across time scales. Fig. 4 shows that, at fast time scales, the most reliable (i.e., with ICC>0.4) FC edges are the ones related to the connectivity between somatomotor and visual regions (Fig. 4A). As the time scale increases, “higher-order” regions start to appear, such as default-mode network and fronto-parietal regions (Fig.4A). Notably, no ICC patterns above 0.4 could be obtained from 100 instances of surrogate data obtained by randomly shuffling subject labels in the dataset (Fig. S3, see also Methods for details on the implementation).

**Fig. 4.**
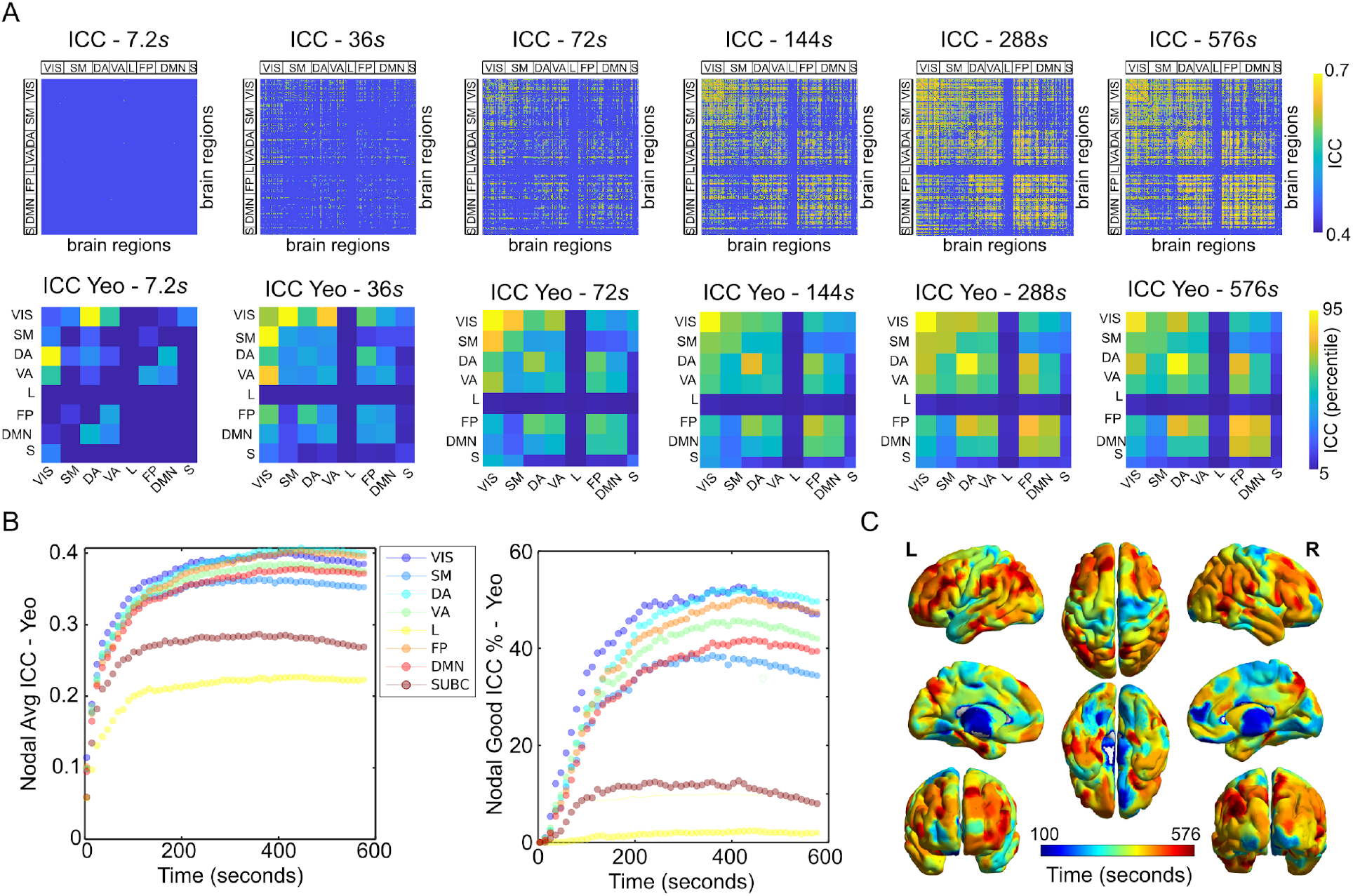
Time scales of brain fingerprints. **A)** Top: Edgewise intra-class correlation (ICC) for the most identifiable frame as function of temporal scale. The ICC matrices are thresholded at 0.4, which is usually a lower limit to define a “good” ICC score ^13,14^. The ICC matrices are ordered according to the seven resting state subnetwork organization proposed by Yeo and colleagues11, specifically: Visual (VIS), somatomotor (SM), dorsal attention (DA), ventral attention (VA), Limbic (L), Frontoparietal (FP) and default mode network (DMN). For completeness, an eight subcortical subnetwork (S) was added at the end (see Brain Atlas in Methods for details). Bottom: the ICC edgewise scores on top are averaged across Yeo functional networks, to better visualize patterns within and between functional subsystems. **B)** The nodal ICC (sum across rows of the ICC matrices) per Yeo functional network is plotted across time, for the un-thresholded (left) and thresholded (right) ICC matrices. **C)** The maximum value across temporal profiles is overlaid onto a brain render, to obtain a brain map of the time scales of human brain fingerprints.

These results reveal a specific time scale for fingerprinting of different functional networks. Indeed, when refining the temporal exploration (see Methods for details), and looking at the nodal counterpart of the ICC profiles across functional networks, one can notice that the temporal fingerprint of each functional subsystem peaks at specific times (Fig. 4B): shorter for subcortical and somatomotor connections, longer for DMN/frontoparietal ones (Fig. 4B, Fig. 4C).

Finally, we looked into temporality of brainprints and the link with behavior. We applied a NeuroSynth meta-analysis based on 50 topic terms onto the brain fingerprint extracted at a specific temporal window, similarly to previous work ^15,16^. We found that brain fingerprints at fast scales are associated with low-order multisensory processing, visual perception, motor/eye movements, as well as affective processing and visuospatial attention (Fig. 5). On the other hand, brain fingerprints at slower time scales are linked to reading comprehension, awareness, verbal semantics, language, social cognition, as well as declarative and working memory (Fig. 5).

**Fig. 5.**
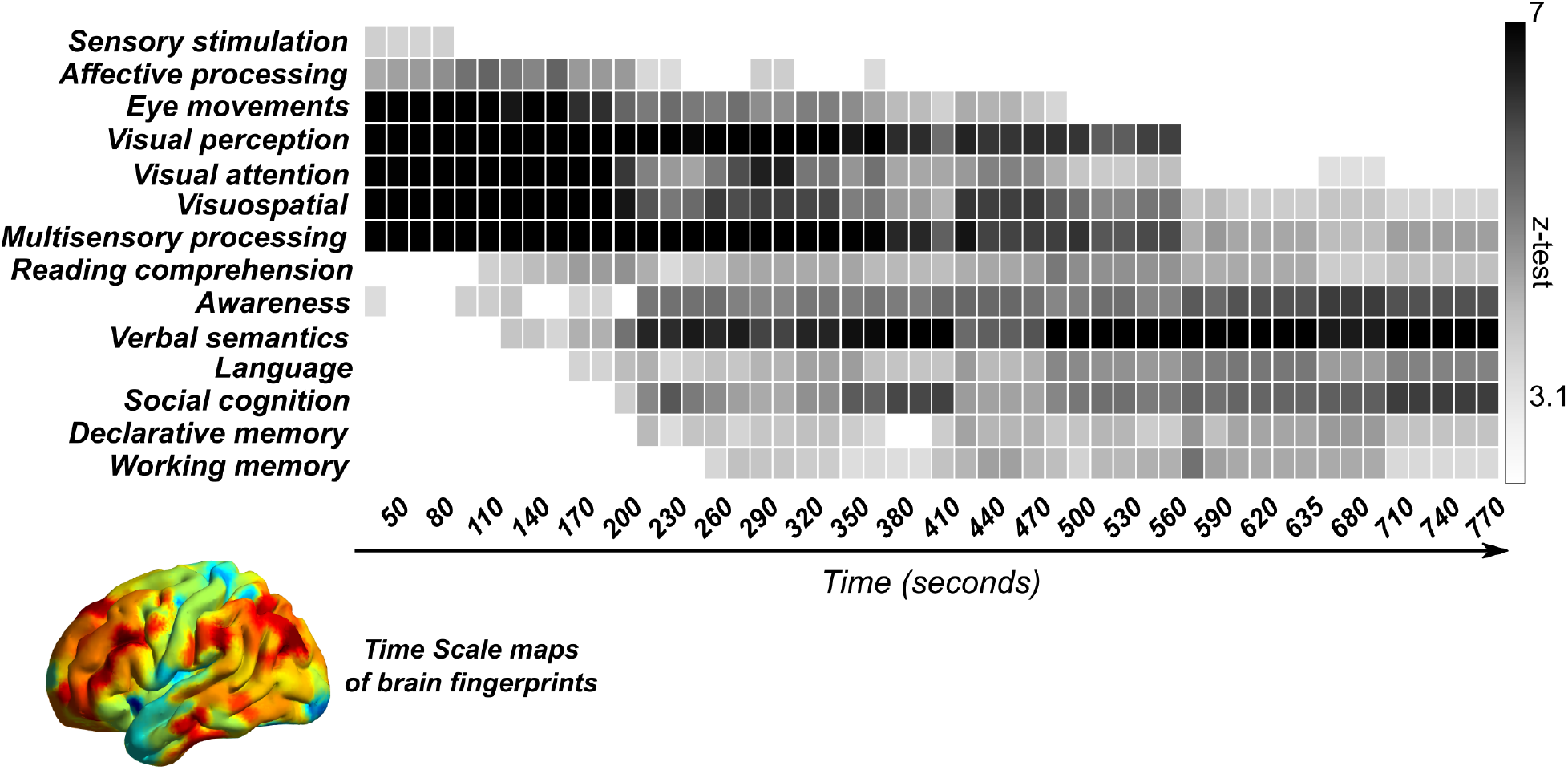
Brain fingerprints associates with behavior across time scales. The NeuroSynth meta-analysis of the brain fingerprints maps across time scales (from 50 seconds to 770 seconds, in steps of 15 seconds) shows a spectrum of association with low-sensory regions at fast time scales, ending up into higher-order processing. The brain fingerprint maps were masked by selecting only the top 25% brain nodes at each time scale.

## Discussion

The neuroscientific community is moving towards open^17^ and reproducible^18^ science, to strengthen and deepen our understanding of the links between cognition^19^, behavior^20^, dysfunction^9^. In this respect, brain fingerprinting has the promise to play a key role providing valuable insights due to its potential inherent to drawing single subject inferences from functional connectivity profiles. Seminal work ^5,10^ showed that brain fingerprints derived from whole resting-state sessions contain uniqueness of each individual functional connectome in the brain areas devoted to “higher-order” cognitive functions, such as frontoparietal and default-mode networks. However, the temporality for identifiability of functional connectomes^10^ has not been addressed before. It relates to the key question: when and at which time scale functional connectomes become unique and thus “identifiable”?

In order to figure out what makes a human brain identifiable at the level of functional neuroimaging correlates, we explored the temporality of brainprints along two complementary directions: 1) *what is the optimal time scale* at which fingerprints integrate sufficient information, 2) *when* does better identification of a fluctuating pattern happen. Using dynamic identifiability (Fig. 1), we showed that the best identification emerges at longer time scales (Fig. 2); however, short transient bursts of “Identifiability” persists even when looking at shorter functional interactions. As a matter of fact, even at faster time scales, very few frames suffice to identify an individual from the others (Fig. 3). These bursts of identifiability might be strongly associated with neuronal activity, as the regularized deconvolution with the hemodynamic response function did not tamper the identification rates obtained (Fig. S1). It is known that mammalian cortical neurons interact in functionally relevant oscillating networks, which span across a broad frequency range ^21^. There is also recent evidence that episodic local field-potential oscillations elicit whole-brain fMRI activity: for instance, hippocampal population burst appears temporally bounded by massive activations of association and primary cortical areas in monkeys ^22^. Based on our findings on human brainprints, we conjecture that this burst of neuronal activity might be one of the sources of this subject-specificity, and therefore closely related to the transient burst in identifiability observed (Fig.3, Fig. S1).

Furthermore, previous studies ^5,10^ showed that the main drivers of the uniqueness of each individual functional connectome reside in the brain areas devoted to “higher-order” cognitive functions, such as frontoparietal and default-mode networks. We found compelling evidence that different parts of connectome fingerprints are optimal at different time scales (Fig. 4). Each region contributes differently to fingerprinting at a specific time scale; i.e., more visual-somatomotor at short temporal windows, more frontoparietal-DMN driven with increasing temporal windows. These findings open up interesting speculation on the link between individual connectivity profiles and the information content associated with the windowed BOLD time series. Maybe higher-order cortices fluctuations contain more slow-long range information that is lost when looking at short windows within them?

As a matter of fact, human brain regions appear to be broadly differentiated into different aspects of behavior and cognition, and the temporal dynamics of neuronal populations across the cortex are thought to be reflective of this specialization ^23–26^. For example, primary sensory neurons are tightly coupled to changes in the environment, firing rapidly to the onset and removal of a stimulus, and showing characteristically short intrinsic timescales. In contrast, neurons in cortical association (or transmodal) regions, exhibit longer firing patterns when a person is engaged in higher-order cognitive tasks ^23^. We here hypothesized that the difference in the brain prints spatial patterns might be tightly linked to the neuronal timescale of the different cognitive processes taking place in a resting human brain (Fig. 4).

Intriguingly, the temporal scales of fingerprinting can be related to behavior in a meaningful way (Fig. 5). Using a meta-analytic approach, we showed that there is a broad spectrum of associations with behavior. At faster time scales, human brain fingerprints are linked to multisensory stimulation, eye movements, affective processing, visuospatial attention. At slower time scales instead, we find higher-cognitive functions, such as language and verbal semantics, awareness, declarative and working memory, social cognition (Fig. 5). This finding reveals for the first time the link between the behavioral relevance of specific functional networks and the associated time scale at which they are manifested.

Notably, these findings are also in line with recent evidence that neuronal timescales follow cytoarchitectonic gradients across the human cortex and are relevant for cognition in both short and long terms ^23^. Particularly, neuronal timescales increase along the principal sensorimotor-to-association axis across the cortex and align with macroscopic gradients of gray matter myelination (T1w/T2w ratio) and synaptic receptor and ion channel gene expression ^23^. Previous work also suggests that functional cortical networks are organized as two large ring-shaped networks (differentiated by their preferred information processing mode) ^27^. The first ring comprises visual, auditory, somatosensory, and motor cortices that process real time world interactions; the second ring, including parietal, temporal, and frontal regions with networks dedicated to cognitive functions, emotions, biological needs, and internally driven rhythms. There is evidence that the patterns of gene expression organize the cortex into two sets of regions that match the two rings ^27^. Overall, the correspondence between the temporal maps brainprints and genetic/cytoarchitectonic profiles, as well as behaviorally-relevant gradients ^16^, opens up more fascinating questions on the relationships between these gradients and human brain identifiability.

This work comes with some limitations. First, the impact of the choice of the brain atlas should be further verified. Second, we examined temporality of brainprints here using sliding window analysis. Future studies should also consider other approaches, such as edgewise connectivity ^28^ or more advanced dynamic functional connectivity models ^29^ . Investigation on the relationship between transient activation and brainprints (Fig. S2) suggests that identification is unlikely to be explained only as a byproduct of hemodynamics. Hence, it would be also interesting to compare the findings from this study, which are based on fMRI resting-state data, with data coming from fMRI task analysis, or even other neuroimaging modalities, such as EEG or MEG. This would allow us to extend the range of accessible time scales across modalities, for dynamic identification. This work opens also the avenue of relating functional brainprints with underlying structural architecture. Recent studies have shown that building FC matrices from (very) long RS fMRI sessions leads to very good proxies for SC ^30^. Similarly, it has been shown that longer fMRI sessions (up to a plateau ^10^) improve identification. Furthermore, function-structure dependency was recently shown to follow a brain pattern extremely consistent with the gradient found here for brainprints ^15^. Future work on brainprints and their association with structural connectivity seems therefore worth exploring.

In sum, we have here explored for the first time the temporality of the human brain fingerprint. We have shown that fingerprints are intertwined with the time scales of functional brain connectivity, and possibly associated with transient bursts in brain activity. This investigation is promising based on these first findings, and represents the first step towards a better understanding of *what* and *when* makes our brains unique.

## Materials and Methods

### HCP data: functional preprocessing

The fMRI dataset used in this work is from the Human Connectome Project (HCP, http://www.humanconnectome.org/), Release Q3. We assessed the 100 unrelated subjects (54 females, 46 males, mean age = 29.1 ± 3.7 years) as provided from the HCP 900 subjects data release ^17,31^ . Per HCP protocol, all subjects gave written informed consent to the Human Connectome Project consortium. The fMRI resting-state runs (HCP filenames: rfMRI_REST1 and rfMRI_REST2) were acquired in separate sessions on two different days, with two different acquisitions (left to right or LR and right to left or RL) per day ^32,33^. For all sessions, data from both the left-right (LR) and right-left (RL) phase-encoding runs were used to calculate connectivity matrices, in order to have four functional connectomes (one LR test-retest pair, one RL) per subject. For this study, we employed the minimally preprocessed HCP resting-state data ^32^, with the following preprocessing steps. First, we applied a standard general linear model (GLM) regression which included: detrending and removal of quadratic trends; removal of motion regressors and their first derivatives; removal of white matter (WM), cerebrospinal fluid (CSF) signals and their first derivatives; global signal regression (and its derivative). Secondly, we bandpass filtered the time series in the range [0.01 0.15] Hz. Finally, the voxelwise fMRI time series were averaged into their corresponding brain nodes of the atlas (see next section, **Brain Atlas**), and then z-scored.

### Brain atlas

We employed a cortical parcellation into 400 brain regions as recently proposed by Schaefer and collaborators^34^ (freely available at ^28^https://github.com/ThomasYeoLab/CBIG/tree/master/stable_projects/brain_parcellation/Schaefer2018_LocalGlobal). For completeness, 16 subcortical regions and 3 cerebellar regions were also added, as provided by the HCP release (filename “Atlas_ROI2.nii. gz”), resulting in a final brain atlas of 419 brain nodes.

### Dynamic Functional Connectivity estimation

To assess brain fingerprints across time scales, we performed sliding window analysis ^29^. The sliding window scheme is the following: first a temporal window, parameterized by its length *w*, is chosen, and within the temporal interval that it spans (i.e., from time *t=1* to time *t=w*), connectivity is computed between each pair of timecourses as Pearson correlation coefficient, producing one instance of the “dynamic functional connectome” (Fig. 1A). Then, the window is shifted by a step *T*, and the same calculations are repeated over the time interval [*t=1 + T*, *t=w + T* ]. This process is iterated until the window spans the end part of the time courses, to obtain a set of connectivity matrices (i.e., dynamic functional connectomes), summarizing the temporal evolution of whole-brain functional connectivity (Fig. 1A). In this work, we started by exploring six different window lengths *w*, specifically of [7.2 36 72 144 288 576] seconds each, and the sets of dynamic functional connectomes associated to them (the number of brain regions is equal to 419). The choice of the shortest window length did not consider the recommendation of previous work^26^ based on stationarity assumptions, instead we opted to allow transient non-stationary events at the level of edgewise FC^28^ to be fully present in the dynamic functional connectomes, at the risk of potential aliasing. The window step *T* was fixed to 7.2 seconds in this study (note that since the TR of HCP is 0.720 *s*, 7.2 *s* corresponds to 10 fMRI data points). Specifically, we studied the evolution of brain fingerprinting across different temporal windows, as detailed in the next section.

### Dynamic Identification

The idea on dynamic identification was inspired by recent work on maximization of connectivity fingerprints in human functional connectomes ^10^. In that work, the authors defined a mathematical object known as “Identifiability matrix”, which is a similarity matrix encoding the information about the self similarity (Iself, main diagonal elements) of each subject with herself/himself, across the test/retest sessions, and the similarity of each subject with the others (or I*others*, off diagonal elements). The similarity between two functional connectomes was quantified with the Pearson’s correlation coefficient between the entries of the connectivity matrices. The difference between I*self* and I*others* (denominated “Differential Identifiability” or “Differential Identification” - Idiff) provides a robust score of the fingerprinting level of a specific dataset ^10^. This idea needs to be extended in the case of dynamic functional connectome evaluation, since in addition to the test/retest set, the set of dynamic “frames” of connectivity are estimated (Fig. S1A, Fig. S1B). For a fixed window length *w*, the resulting “Dynamic Identifiability matrix” (Fig. S1C) *d*I is then a block-diagonal matrix, where each block represents the self-similarity within the dynamic functional connectome frames of a specific subject. The off-diagonal blocks, in this representation, encode instead the between-dFC frames similarity across different subjects (dynamic I*others*). Let 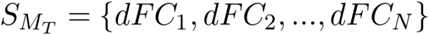 be the set of dFC frames in the test session, for a specific subject M. Similarly, let 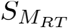 represent the set of dFC frames in the retest session, for the same subject M. We can then define the dynamic I*self (d*I*self)* for subject M as:

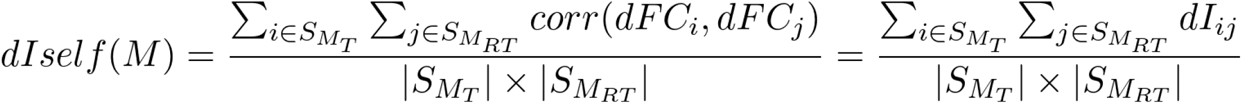

Where 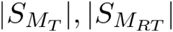 define the cardinalities of the sets. Similarly, let 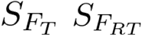 define the sets for a different subject F. We can define dynamic I*others* as:

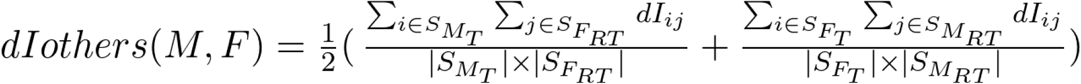

And hence:

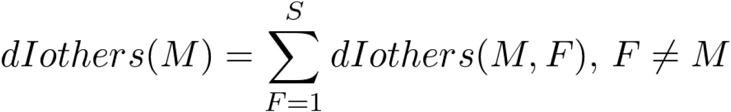

Where the summation is over the total number of subjects S other than M. Finally, dynamic differential identifiability for a subject M results in:

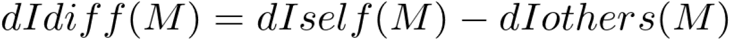

And the average *d*Idiff:

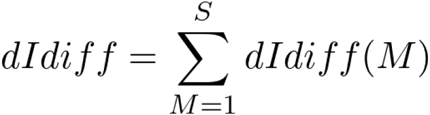

We use dynamic differential Identifiability to explore connectivity fingerprints across different window lengths. Note that we first evaluated dynamic identification of the LR and RL connectome pairs separately, and then averaged the corresponding LR/RL Dynamic Identifiability matrices into one.

### Maximum dynamic dFC frame selection

The dynamic identification framework described above provides the average behavior of fingerprinting within and between dynamic functional connectome frames. However, there might be dynamic FC frames where identification is higher than others, which might not be captured by the average behavior depicted by *d*Idiff. To cover the necessity of that, for each subject we sorted the dFC frames in test retest according to their similarity, from highest to lowest, based on their *d*Iself_ij_ value (see *d*Iself equation above). We then explored *d*Iself, *d*Iothers and *d*Idiff when iteratively adding dFC frames one at the time, starting from the best matching ones, and then proceeding based on their similarity values.

### Transient (total activation) analysis

The TA framework^11^ incorporates two main features for fMRI data processing: (1) each voxel’s BOLD time course is deconvolved from the temporal blur introduced by the hemodynamic response, leading to the activity-inducing signal that is supposed to show block-type activation patterns (without any prior knowledge on the timing and duration of these blocks); (2) BOLD signals should show a spatial smoothness, which is supposed to be stronger within anatomical atlas regions than across. With that aim, TA solves a convex optimization problem that consists of a least-squares data-fitting term combined with spatial and temporal regularization terms. TA produces de-noised and well-behaving reconstructions of the activity-related, activity-inducing signals, decoupled from the hemodynamics. We used this framework to study dynamic identification properties of dFC obtained from transient brain activity, and compare it to the results obtained with the original dFC frames.

### Dynamic edgewise identification

We quantified the edgewise reliability of individual dynamic FC frames across different temporal windows by evaluating the intraclass correlation coefficient^36^, similarly to previous work^10^. ICC is a widely used statistical measure to assess the percent of agreement between units of different groups. It describes how strongly units in the same group resemble each other. The stronger the agreement, the higher its ICC value. We used ICC to quantify to which extent the connectivity value of an edge in an FC frame (i.e., FC value between two brain regions) was consistent across test/retest acquisitions and could separate within- and between-subject data. In other words, the higher the ICC, the higher the “fingerprint” of the edge connectivity. Note that we thresholded the resulting ICC matrices at 0.4, which is usually a lower limit to define a “good” ICC score ^13,14^. Finally, the nodal strength of the ICC edgewise matrix (i.e., sum over columns, evaluated with and without thresholding the ICC matrix, see also Fig. 4) was used as a “nodal fingerprinting score” of how central each brain region is to connectome identification.

### Significance of dynamic edgewise identification

In order to better characterize the ICC results in dynamic functional connectomes presented in Figure 4, we performed a permutation testing analysis. Concretely, we evaluated ICC scores in 100 surrogate datasets where subject labels have been randomly shuffled (Fig. S3). Comparing these ICC scores to the original ICC scores presented in Figure 4A allowed us to evaluate the extent to which the best matching connectivity patterns in test and retest datasets at different time scales are unique to the subjects.

### Brain fingerprints and behavior

A NeuroSynth meta-analysis [https://neurosynth.org/] similar to the one implemented in previous studies ^15,16^ was conducted to assess topic terms associated with brainprints across time scales. 50 binary masks of brain fingerprints at different time scales (from 50 s windows to 770 s, in steps of 15 s) were obtained by selecting the top 25 percentile of ICC nodal strength of each brain map, and served as input for the meta-analysis, based on 50 topic terms. Terms were ordered according to the weighted mean of the resulting z-statistics for visualization.

## Acknowledgments

Data were provided [in part] by the Human Connectome Project, WU-Minn Consortium (Principal Investigators: David Van Essen and Kamil Ugurbil; 1U54MH091657) funded by the 16 NIH Institutes and Centers that support the NIH Blueprint for Neuroscience Research; and by the McDonnell Center for Systems Neuroscience at Washington University. EA acknowledges financial support from the SNSF Ambizione project "Fingerprinting the brain: network science to extract features of cognition, behavior and dysfunction“ (grant number PZ00P2_185716). MGP was supported by the CIBM Center for Biomedical Imaging, a Swiss research center of excellence founded and supported by Lausanne University Hospital (CHUV), University of Lausanne (UNIL), Ecole polytechnique fédérale de Lausanne (EPFL), University of Geneva (UNIGE) and Geneva University Hospitals (HUG). We would like to thank Dr. Emanuela De Falco for insightful discussions.

## Code availability

The code (in MATLAB) used for this analysis will be made available upon acceptance of the manuscript on EA EPFL webpage and a git repository.

## Author Contributions

EA. MGP and RL processed the data; DVD and EA conceptualized the study; EA designed the framework and performed the connectivity analyses; EA and YF performed the activity inducing analysis; all authors interpreted the results and wrote the manuscript.

