## Supplementary Information for "*When* makes you unique: temporality of the human brain fingerprint"

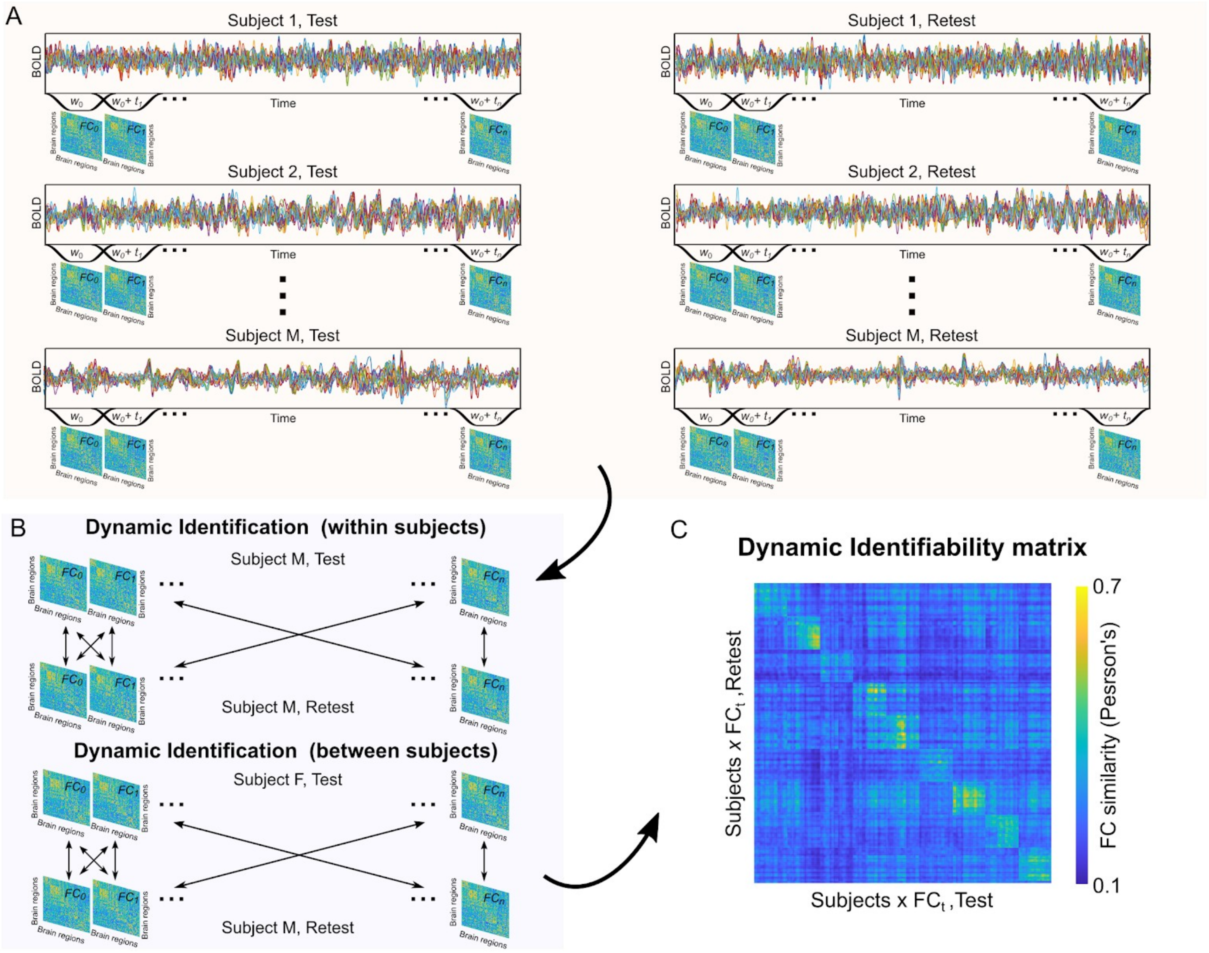


**Fig. S1 Dynamic Identifiability scheme.** Workflow scheme for dynamic fingerprints identification. **A)** First, sliding window functional connectomes (dFC) are computed for each individual BOLD time course, for test and retest sessions. **B)** The similarity between dFC frames (within and between subjects), across test and retest sessions, is then computed. **C)** Finally, the Dynamic Identifiability matrix is computed. This object can be used to visualize and summarize the evolution of brain connectivity fingerprints across temporal scales (i.e., dFC windows).

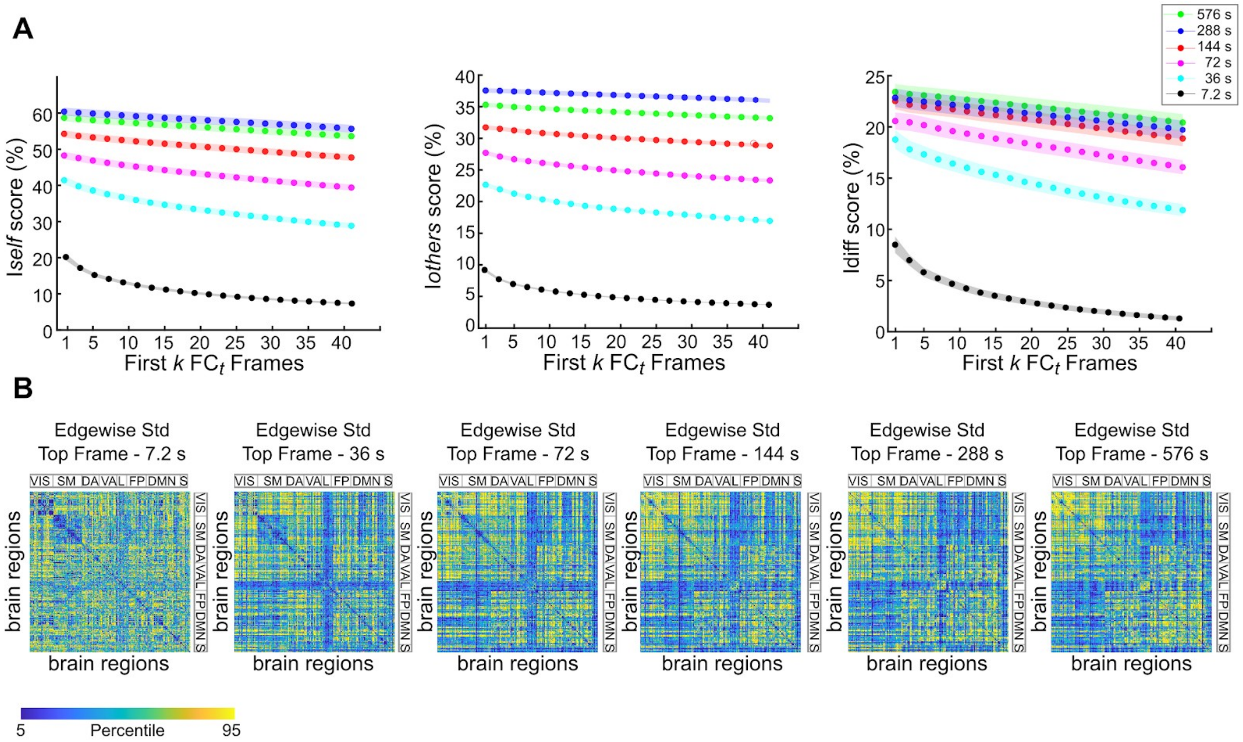


**Fig. S2 Brain fingerprint frames and neuronal activity. A)** Evaluation of brain fingerprints across temporal scales, when ranking dFC frames based on individual *d*Iself, in descending order. In this case, the fingerprints are computed on the deconvoled transient signals (activity inducing, [11]), in the attempt to capture transient neuronal activity from BOLD signals associated with individual identification. **B)** Edgewise standard deviation of the best matching "activity inducing" dFC frames, across temporal scales. The matrices are ordered according to the seven resting state subnetwork organization proposed by Yeo and colleagues[12], specifically: Visual (VIS), somatomotor (SM), dorsal attention (DA), ventral attention (VA), Limbic (L), Frontoparietal (FP) and default mode network (DMN). For completeness, an eight subcortical subnetwork (S) was added at the end (see Brain Atlas in Methods for details).

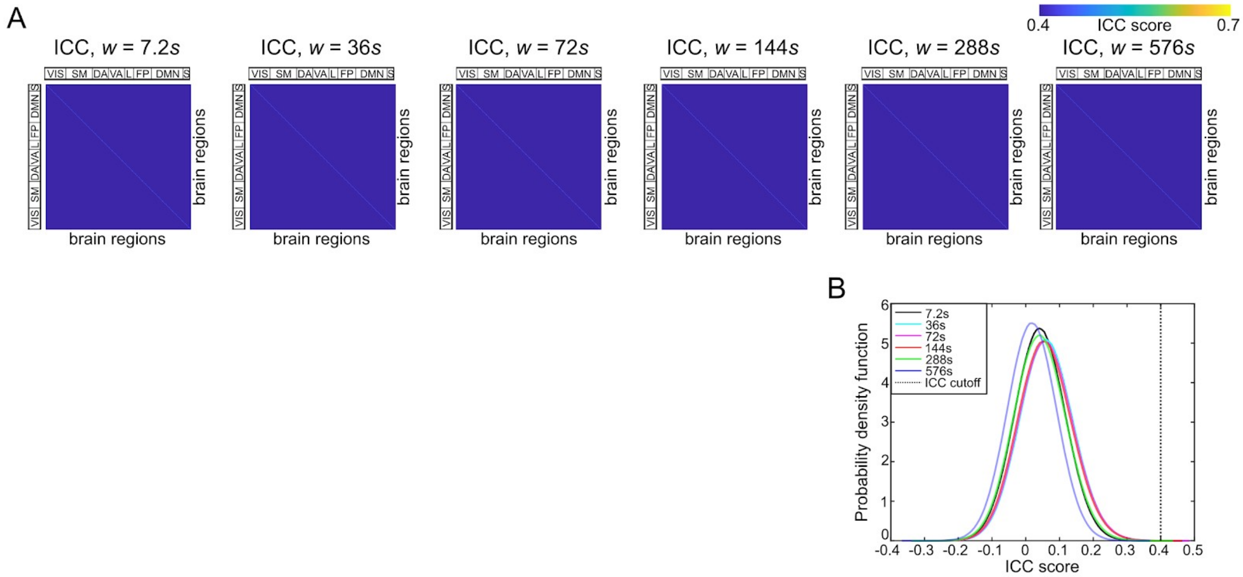


**Fig. S3 Significance of dynamic brain fingerprints. A)** Edgewise intra-class correlation (ICC) for the most identifiable frame, across temporal scales, averaged across 100 runs obtained after randomly shuffling the subjects (see Methods for details). The ICC matrices are thresholded at 0.4, which is usually a lower limit to define a “good” ICC score [13,14]. **B)** Probability density function of the (unthresholded) ICC scores for the random shuffling across the 100 runs. Note how the random shuffling distributions are upper bounded by the good ICC cutoff of 0.4.
